# Characterization of SARS-CoV-2 N protein reveals multiple functional consequences of the C-terminal domain

**DOI:** 10.1101/2020.11.30.404905

**Authors:** Chao Wu, Abraham J. Qavi, Asmaa Hachim, Niloufar Kavian, Aidan R. Cole, Austin B. Moyle, Nicole D. Wagner, Joyce Sweeney-Gibbons, Henry W. Rohrs, Michael L. Gross, J. S. Malik Peiris, Christopher F. Basler, Christopher W. Farnsworth, Sophie A. Valkenburg, Gaya K. Amarasinghe, Daisy W. Leung

## Abstract

Nucleocapsid protein (N) is the most abundant viral protein encoded by SARS-CoV-2, the causative agent of COVID-19. N plays key roles at different steps in the replication cycle and is used as a serological marker of infection. Here we characterize the biochemical properties of SARS-CoV-2 N. We define the N domains important for oligomerization and RNA binding that are associated with spherical droplet formation and suggest that N accessibility and assembly may be regulated by phosphorylation. We also map the RNA binding interface using hydrogen-deuterium exchange mass spectrometry. Finally, we find that the N protein C-terminal domain is the most immunogenic by sensitivity, based upon antibody binding to COVID-19 patient samples from the US and Hong Kong. Together, these findings uncover domain-specific insights into the significance of SARS-CoV-2 N and highlight the diagnostic value of using N domains as highly specific and sensitive markers of COVID-19.

## Introduction

Severe acute respiratory syndrome coronavirus 2 (SARS-CoV-2) is a novel coronavirus and the causative agent of COVID-19. Coronavirus has a single-stranded, positive-sense RNA genome. The genome encodes four major structural proteins: spike (S), envelope (E), membrane (M), and nucleocapsid (N). The N protein is the second most proximal to the 3’ end of the genome and is one of the most abundantly expressed viral proteins given the multifunctional roles N has during viral replication and assembly (Fung and Liu, 2019; Kim et al., 2020; McBride et al., 2014; Perlman and Netland, 2009). It is estimated that 1,000 copies of N are incorporated into each virion versus only 100 copies of S (Bar-On et al., 2020). Whereas N exists mostly in a phosphorylated state in the cytoplasm, it is predominantly in a dephosphorylated state in mature virions, suggesting that genome packaging is regulated by the phosphorylation state of N (Wu et al., 2014; Wu et al., 2009).

A critical function of N is to encapsidate the ssRNA viral genome to evade immune detection and to protect the viral RNA from degradation by host factors (Chang et al., 2014; McBride et al., 2014). N has two structural domains (**Figure 1A**): a N-terminal domain (N_NTD_; amino acid residues 44-176) and a C-terminal domain (NCTD; residues 248-369). N_NTD_ is often referred to as the RNA-binding domain (RBD), although the exact regions involved in RNA binding are not yet well defined (Chang et al., 2014; Grossoehme et al., 2009; Gui et al., 2017; Kang et al., 2020; McBride et al., 2014). The isolated N_CTD_ exists as a dimer in solution and is potentially involved in RNA binding as well (Bouhaddou et al., 2020; Gui et al., 2017; Takeda et al., 2008). A conserved serine/arginine rich-linker region (N_LKR_) connects the N_NTD_ and the N_CTD_. Phosphorylation of residues in the LKR is believed to regulate discontinuous transcription, particularly for shorter subgenomic mRNA closer to the 3’ end during early stages of replication (Wu et al., 2014; Wu et al., 2009). The LKR, along with Narm and Carm, have been shown to be intrinsically disordered (Chang et al., 2014; Cubuk et al., 2020). However, the LKR is similarly conserved as the NTD and CTD (**Figure 1B** and **Supp. Figure 1B**), supporting its essential functional role across strains of coronaviruses (Chang et al., 2014), while the Narm and Carm are the least conserved regions.

**Figure 1.**
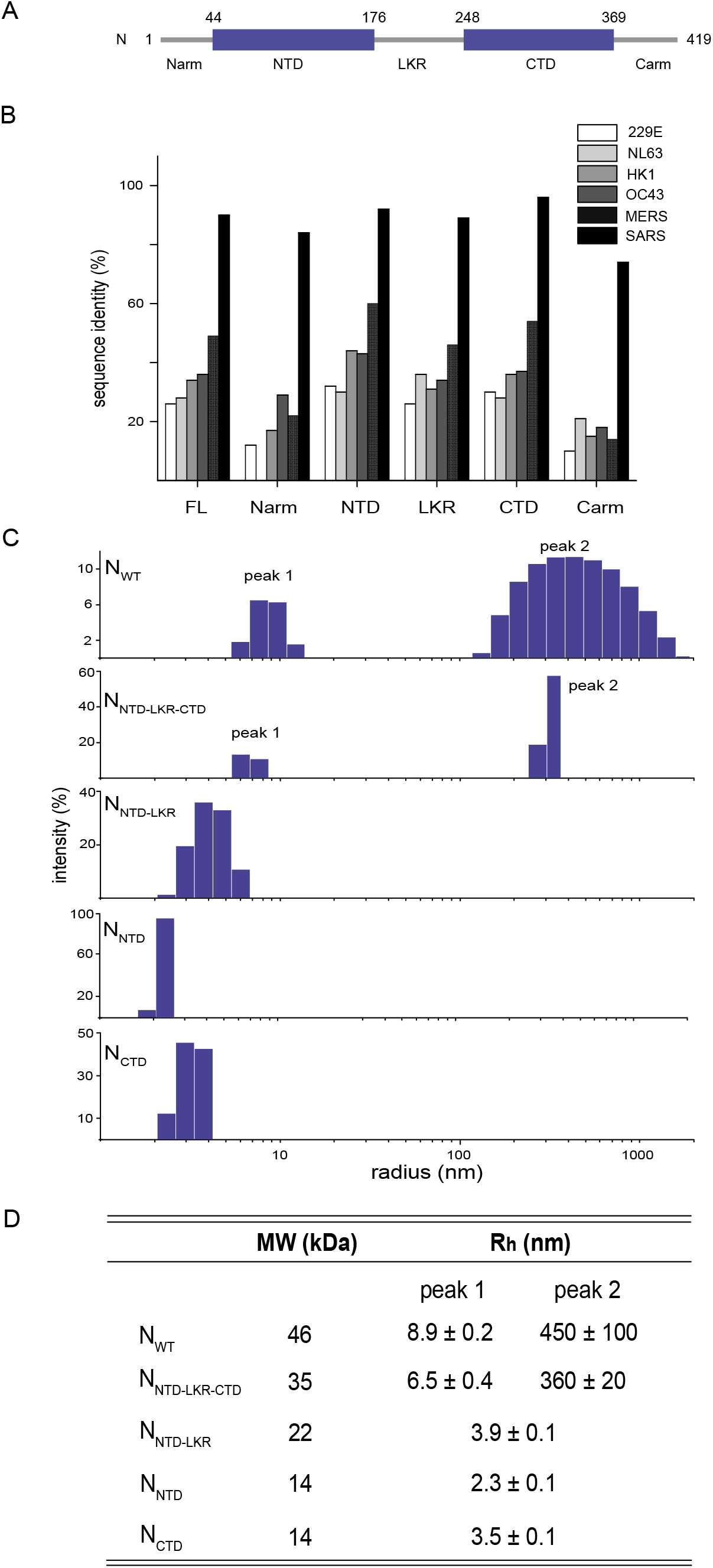
Characterization of N oligomerization with dynamic light scattering. **A**. Domain architecture of N. N has two structural domains: NTD and CTD. The sequence between NTD and CTD is a linker region (LKR) containing a serine-arginine rich motif. The Narm, LKR, and Carm are predicted to be disordered based upon sequence analysis. **B.** Sequence identity between N of SARS-CoV-2 and common cold coronaviruses and other epidemic severe coronaviruses MERS-CoV and SARS-CoV. **C.** Measurements of N oligomerization with dynamic light scattering. Measured hydrodynamic radii, Rh, are reported in D. **D.** Table summarizes the DLS data for all constructs. Numbers are reported as average and standard deviation of three experiments.

Given its abundant expression and conservation within the genome, N has been used as an antigen for serology tests (Chew et al., 2020; Tang et al., 2020a, b), such as the widely used Roche Elecsys Anti-SARS-CoV-2 assay and Abbott SARS-CoV-2 IgG assay (Tang et al., 2020a, b). Comparison of the serological response to the entire proteome of the SARS-CoV-2 virus using a luciferase immunoprecipitation assay (LIPS) (Hachim et al., 2020) showed that N-specific antibodies dominated the overall antibody response. Furthermore, the T cell responses directed towards N are highly immunodominant in SARS-CoV and SARS-CoV-2 infection, with N-specific memory T cell responses evident 17 years after the initial SARS-CoV infection (Le Bert et al., 2020). Due to epitope conservation of the N protein, there is some T cell cross-reactivity towards the N of the SARS-CoV-2 (Le Bert et al., 2020). The N protein stability, RNA binding characteristics, abundance, and conservation altogether impact immunogenicity for T and B cell immunities.

Previous studies, including our own, have examined fundamental properties of nucleocapsid or nucleoproteins (N) from RNA viruses, which revealed overlapping and unique functions (Arragain et al., 2019; Ding et al., 2016; Lu et al., 2020; Luo et al., 2020; Raymond et al., 2010; Su et al., 2018; Wan et al., 2017). These include the insights into RNA binding, oligomerization, as well as potential roles in RNA synthesis and immune evasion. However, such studies for SARS-CoV-2 N protein remain incomplete. To address this gap, here we use a series of biochemical and biophysical assays to probe essential functions of N. Our results reveal that oligomeric N provides a continuous RNA binding platform and that N-RNA has different morphologies, including spherical droplets. Our data also suggest how phosphorylation could modulate these processes, which can be exploited by drugs targeting of these cellular processes (Bouhaddou et al., 2020). Given the recent progress in SARS-CoV-2 spike-based vaccines, knowledge of N will likely provide the basis to differentiate individuals with immunity due to natural infections from those that are immunized. Such tools will play a critical role in managing herd immunity. In addition, our domain-specific insights into the immunogenicity of N provide opportunities to further enhance sensitivity and specificity of serology testing. Therefore, insights gained here regarding N protein function and regulation are critical for improving diagnostic testing for the ongoing COVID-19 pandemic and future outbreaks.

## Results

### Multiple domains within N protein are important for oligomerization

To characterize how the domains of N contribute to oligomerization, we used dynamic light scattering (DLS) to determine the hydrodynamic properties of isolated N domains. The results reveal that there are two major oligomeric species for wildtype (WT) N (N_WT_; 46 kDa), one with a hydrodynamic radius (R_h_) of 8.9 nm and another of 450 nm (**Figure 1C and 1D**). For comparison, the R_h_ values for maltose binding protein (44 kDa) and bovine serum albumin (66 kDa) are 2.9 nm and 3.7 nm, respectively. Removal of the Narm and Carm (N_NTD-LKR-CTD_) generates two major species, similar to N_WT_. However, both N_NTD-LKR-CTD_ populations display reduced polydispersity (peak width, **Supp. Figure 2A**) in the high order oligomeric species, suggesting that both arms contribute to N oligomerization. Removal of the CTD (N_NTD-LKR_) results in a single peak representing a dimeric species (R_h_ = 3.9 nm), but with considerable polydispersity. N_NTD_ and N_CTD_ alone are stable domains; N_NTD_ is a monomer (R_h_ = 2.3 nm) whereas N_CTD_ is a dimer in solution (R_h_ = 3.5 nm). Exact mass measurement by denaturing mass spectrometry yields values corresponding to the expected sequence (± 1 Da) and supports the identity of the constructs used here (**Supp. Figure 2B-E**).

### N provides an oligomerization platform for high affinity RNA binding

N encapsidates viral genomic ssRNA in a sequence-independent manner (Chang et al., 2014). To gain insight into how each N domain contributes to ssRNA binding, we measured fluorescence polarization (FP) upon binding of a FITC-labeled 20-nt ssRNA (Sequence: UUUCACCUCCCUUUCAGUUU) (**Figure 2A**). We find that NWT binds the 20-nt ssRNA with high affinity (*K_D_* = 0.007 ± 0.001 μM). Removal of the Narm and Carm do not impact ssRNA binding (*K_D_* = 0.006 ± 0.002 and 0.006 ± 0.002 μM for N_NTD-LKR-CTD-_ Carm and N_NTD-LKR-CTD_, respectively) (**Figure 2B**). The individual domains N_NTD_ and N_CTD_ have low affinity binding (*K_D_* = 20 ± 10 and 13 ± 5 μM, respectively), both of which bind with increased affinity after inclusion of the LKR region (**Figure 2B–2D**). Interestingly, addition of CTD that dimerizes onto NTD-LKR improves the binding affinity to the single digit nM range (**Figure 2C**). The affinity increase does not happen when CTD is provided *in trans* (compare N_NTD-LKR_ + N_CTD_ with N_NTD-LKR-CTD_). This affinity increase also occurs when NTD is added to LKR-CTD *in cis* (**Figure 2D**, compare N_LKR-CTD_ + N_NTD_ with N_NTD-LKR-CTD_). Similar binding curves and *K_D_* values were obtained when fluorescence anisotropy values were converted from polarization (**Supp. Figure 3A**). Collectively, our data show that NTD, CTD, and LKR all contribute to ssRNA binding, and the presence of three domains in tandem confers N with high affinity binding.

**Figure 2.**
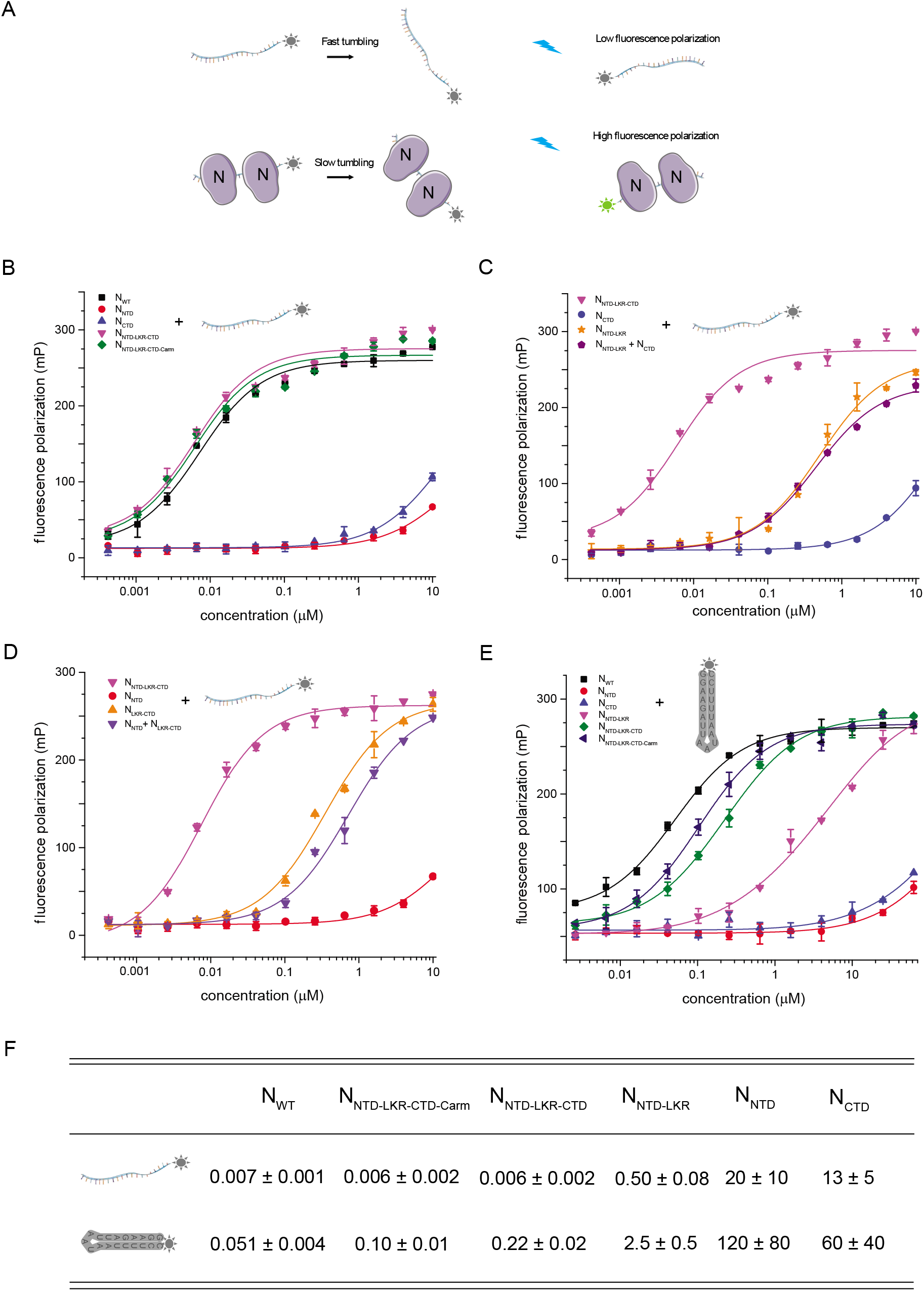
SARS-CoV-2 nucleocapsid protein binds ssRNA with high affinity. **A.** Principles of fluorescence polarization assay to measure RNA binding. Increasing concentrations of N was titrated into 1nM of labeled RNA. Protein binding to labeled RNA leads to slower tumbling of labeled RNA, resulting in increased fluorescence polarization. **B.** Fluorescence polarization binding curves of N constructs to a 20-nt ssRNA. The fitted K_D_ values are 0.007 ± 0.001 μM (N_WT_, black square), 16 ± 12 μM (N_NTD_, red circle), 13 ± 5 μM (N_CTD_, blue up triangle), 0.006 ± 0.002 μM (N_NTD-LKR-CTD_, magenta down triangle), and 0.006 ± 0.002 μM (N_NTD-LKR-CTD-Carm_, green diamond). **C.** Fluorescence polarization binding curves of N constructs to a 20-nt ssRNA. The fitted K_D_ values are 0.006 ± 0.002 nM (N_NTD-LKR-CTD_, magenta down triangle), 13 ± 5 μM (N_CTD_, blue circle), 0.50 ± 0.08 μM (N_NTD-LKR_, orange star), and 0.44 ± 0.04 μM (N_NTD-LKR_ + N_CTD_, purple pentagon). **D.** Fluorescence polarization binding curves of N constructs to a 20-nt ssRNA. The fitted K_D_ values are 0.006 ± 0.002 μM (N_NTD-LKR-CTD_, magenta down triangle), 16 ± 12 μM (N_NTD_, red circle), 0.35 ± 0.04 μM (N_LKR-CTD_, orange up triangle), and 0.72 ± 0.09 μM (N_NTD_ + N_LKR-CTD_, purple down triangle). **E.** Fluorescence polarization binding curves of N constructs to a slRNA. The fitted K_D_ values are 0.051 ± 0.004 μM (N_WT_, black square), 124 ± 84 μM (N_NTD_, red circle), 65 ± 44 μM (N_CTD_, blue up triangle), 2.5 ± 0.5 μM (N_NTD-LKR_, magenta down triangle), 0.22 ± 0.02 μM (N_NTD-LKR-CTD_, green diamond), and 0.10 ± 0.01 μM (N_NTD-LKR-CTD-Carm_, navy left triangle). **F**. Table summarizes K_D_ values (μM) for key constructs binding to ssRNA and slRNA. Numbers are reported as average and standard deviation of two experiments.

N also binds dsRNA and is proposed to disrupt dsRNA structures formed by transcription regulatory sequences (TRS) during discontinuous transcription (Grossoehme et al., 2009; Keane et al., 2012; Sola et al., 2015). We next tested N binding to a stem-loop RNA (slRNA) (Sequence: GGAAGAUUAAUAAUUUUCC) (**Figure 2E**). We find that N_WT_ binds with relatively high affinity (*K_D_* = 0.051 ± 0.004 μM) whereas both N_NTD_ and N_CTD_ alone have very weak binding affinities (*K_D_* = 120 ± 80 and 60 ± 40 μM, respectively). The addition of LKR significantly improves binding, consistent with binding results for ssRNA. Overall, binding of N to slRNA appears similar to binding to ssRNA, but at an order of magnitude lower than that of ssRNA (**Supp. Figure 3B**). This may be due, in part, to the energetic penalty of unfolding the stem loop structure. Furthermore, it seems that Narm and Carm contribute more to slRNA binding than ssRNA because the reduction of *K_D_* for slRNA are more pronounced after removing the Narm and Carm (**Supp. Figure 3C**).

### N-RNA forms spherical droplets, requiring multiple N domains

Purification of recombinant N protein on size exclusion chromatography revealed three populations, including two RNA-bound states (p1 and p2) and an RNA-free state (p3) despite purification in high salt (500 mM) to eliminate RNA binding (**Figure 3A**). Truncation of the Narm results in an increase of the RNA-free peak (p3), suggesting that N truncations can alter the structure of N and correspondingly impact N-related properties, such as RNA binding and oligomerization, thereby potentially impacting the physiological function of N. We observe an even greater shift to p3 when both Narm and Carm were removed, suggesting that both arms contribute to RNA binding interactions. To gain additional insight into the two RNA-bound populations p1 and p2, we visualized these samples by using negative-stain electron microscopy (EM) in near-physiological salt concentrations (150 mM). N_WT_ p1 contains N-RNA with a loose-coil appearance (**Figure 3B, top left**), similar to that observed for other RNA-bound nucleocapsids (Bharat et al., 2012; Mavrakis et al., 2002). Other recent studies have also observed that N protein undergoes liquidliquid phase separation (LLPS) when mixed with RNA(Carlson et al., 2020; Cubuk et al., 2020; Iserman et al., 2020; Jack et al., 2020; Savastano et al., 2020). In agreement with these results, for N_WT_ p2, we mostly observe spheres corresponding to liquid droplets separated from the surrounding buffer (**Figure 3B, top right**).

**Figure 3.**
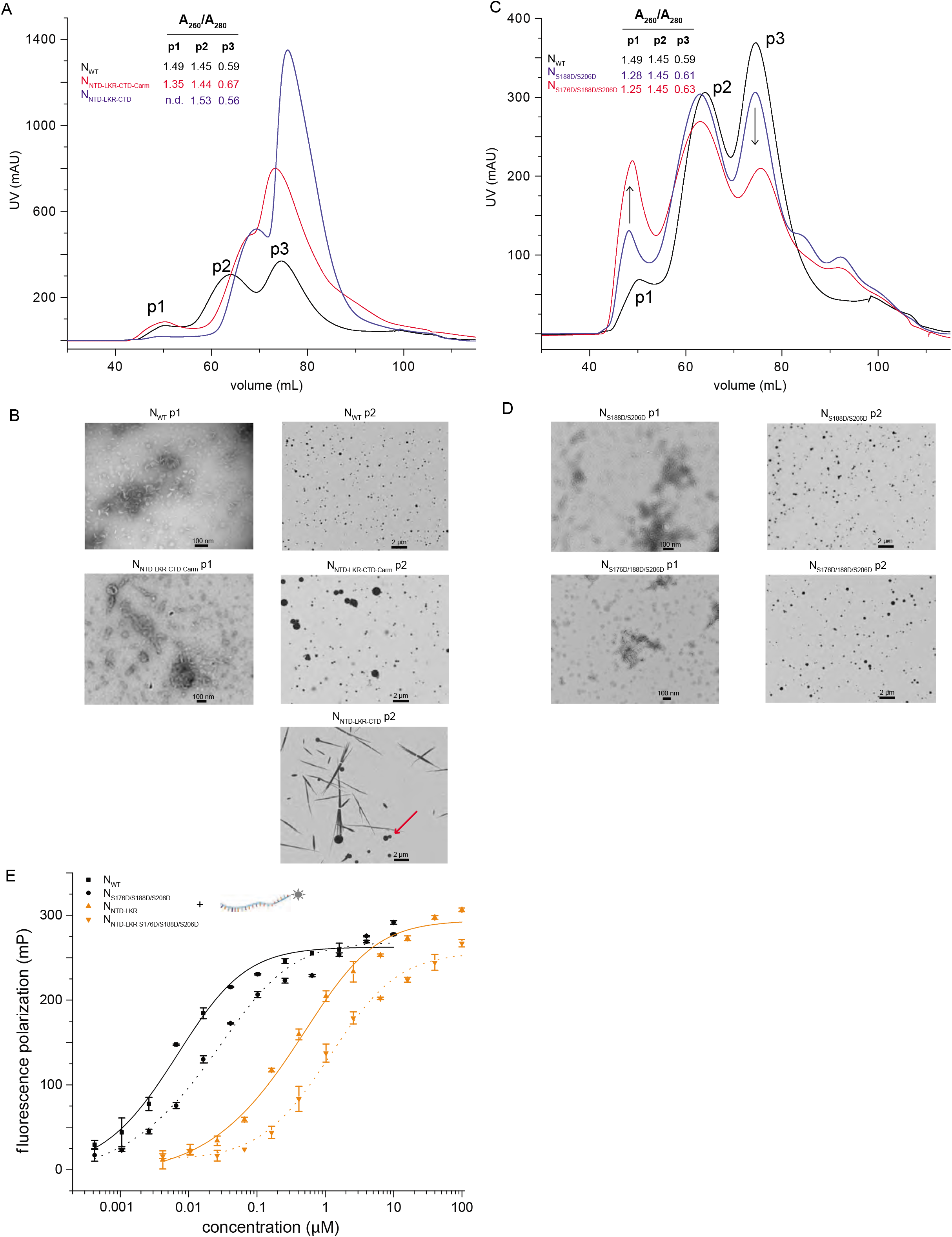
N-RNA phase separates and phosphorylation modulates N-RNA. **A.** Size exclusion chromatography of N constructs (N_WT_, black; N_NTD-LKR-CTD-Carm_, red; N_NTD-LKR-CTD_, blue) in 25 mM HEPES, 500 mM NaCl, 2 mM TCEP, 5% glycerol from a 2 L pellet. Samples from peak 1 (p1) and p2 contain RNA whereas p3 are RNA-free based upon absorbance from the 260/280 ratio. **B.** Negative stain electron microscopy (EM) image of p1 and p2 for NWT, N_NTD-LKR-CTD-Carm_, and N_NTD-LKR-CTD_ in 150 mM NaCl. Samples were diluted into 150 mM NaCl before negative-staining fixation by uranyl acetate. **C.** Size exclusion chromatography of N constructs (N_WT_, black; N_S188D/S206D_, blue; NS1 76D/S188D/S206D, red) in 25 mM HEPES, 500 mM NaCl, 2 mM TCEP, 5% glycerol. **D**. Negative stain electron microscope image of N_S188D/S206D_ and N_S176D/S188D/S206D_ in 150 mM NaCl. **E.** Fluorescence polarization binding curves of N mutants to a 20-nt ssRNA. The fitted K_D_ values are 0.007 ± 0.001 μM (N_WT_, black square), 0.015 ± 0.002 μM (NS1 76D/S188D/S206D, black circle), 0.505 ± 0.075 μM (N_NTD-LKR_, orange up triangle), and 1.1 ± 0.2 μM (N_NTD-LKR S176D/S188D/S206D_, orange down triangle). Numbers are reported as average and standard deviation of two experiments.

We next examined the role of Narm and Carm on high-order assembly. N_NTD-LKR-CTD-Carm_ behaves similarly to N_WT_ and has loose coils in p1 (**Figure 3B, middle left**) and forms spherical liquid droplets in p2 (**Figure 3B, middle right**). However, examination of p2 from N_NTD-LKR-CTD_ (**Figure 3B, bottom right**) revealed a much smaller population of liquid droplets (red arrow) and mostly crystal-like needle aggregates, suggesting that the Carm is important for droplet formation. A transition from spherical liquids to needle-like solids is consistent with the liquid-to-solid transitions observed for other proteins that undergo phase separation(Patel et al., 2015).

### Phosphorylation of LKR modulates RNA binding and higher-order assembly

Previous studies have shown that N is dephosphorylated in mature virions of a closely related strain, SARS-CoV, and predominantly phosphorylated during active transcription and replication in the cytoplasm (Wu et al., 2014). For SARS-CoV, glycogen synthase kinase (GSK)-3 was shown to phosphorylate N at Ser177 (corresponding to Ser176 in SARS-CoV-2 N) (Wu et al., 2009). Phosphorylation of Ser177 is preceded by phosphorylation of Ser189 and Ser207 (Ser188 and 206 in SARS-CoV-2 N) by other priming kinases (Wu et al., 2009). Moreover, N protein phosphorylation has been qualitatively shown to modulate both RNA binding and phase separation(Carlson et al., 2020; Lu et al., 2020; Savastano et al., 2020). To test if phosphorylation impacts RNA binding and the phase separation of SARS CoV-2 N-RNA, we generated a set of N phosphomimics by mutating specific serine residues to aspartate residues. Size exclusion chromatography showed that, compared to N_WT, NS188D/S206D_ produced a reduced RNA-free peak (p3) and an increased RNA-bound peak (p1) (**Figure 3C**). Introduction of S176D to generate N_S176D/S188D/S206D_ resulted in an even greater shift in p1 and p3 distributions, showing how phosphorylation can affect N interactions with RNA. The height of p2 remains relatively the same for all preparations. Examination of these protein peaks using electron microscopy revealed that N_S188D/S206D_ displays similar loose coils in p1 (**Figure 3D, top left**) and spherical droplets in p2 (**Figure 3D, top right**) for the RNA-bound species. Similar observations were made for N_S176D/S188D/S206D_ (**Figure 3D**). To describe this interaction further, we measured ssRNA binding to the N phosphomimics (**Figure 3E** and **Supp. Figure 4A**). N_S176D/S188D/S206D_ displays ~5-fold lower binding affinity to ssRNA compared to N_WT_ binding, a result consistent with previous work examining the impact of LKR phosphorylation on RNA binding(Savastano et al., 2020). We observed a similar trend for the N_NTD-LKR_ construct. Furthermore, binding to slRNA is also affected by these mutations (**Supp. Figure 4B**). Collectively, our data suggest that phosphorylation of the LKR region modulates N interactions with RNA, causing changes in solution properties. Interestingly, there are 14 serine residues in the LKR, of which 13 are found in SARS-CoV, and an increase in phosphorylation in this region may further enhance these changes for RNA interaction and subsequent viral replication and immunogenicity.

### Hydrogen-deuterium exchange mass spectrometry (HDX-MS) locates the RNA binding interface of NTD-LKR

We performed HDX-MS to locate regions that become protected upon RNA binding. Using N_NTD-LKR-CTD_, we observed phase-separation and aggregation upon RNA binding, causing a 100-fold mass spectrometric signal loss for the bound state possibly attributable to phase separation or aggregation inducing poor digestion. However, this problem does not pertain to N_NTD-LKR S176D/S188D/S206D_ where digestion yielded 152 peptides after sequential FXIII and peptic digestion and provided 93.3% sequence coverage (**Supp. Figure 5A**).

HDX analysis of N_NTD-LKR S176D/S188D/S206D_ shows clear protection in four distinct regions of upon RNA binding (aa residues 41-63, 105-108, 146-171, and 213-230) (**Figure 4A** and **4B**). Residues 133-143 are not perturbed by RNA binding, but peptides covering 146-171 show clear protection. The largest differences in HDX are observed where 50-80% of the residues of unbound peptides undergo a burst phase of HDX in the first 10 s (146-156, 163-171, and 213-230 aa), i.e., the peptides cover regions of little hydrogen bonding in the unbound state. When bound to RNA, the fraction of residues participating in the burst phase decreases, resulting in observed protection. Then, HDX either converges over time (146-156, 163-171, and 219-223 aa) consistent with protein conformation or RNA binding dynamics, or the HDX never converges in the timescale of the experiment (222-230 aa), consistent with relatively static binding. Interestingly, peptides covering 103-108 aa and 156-159 aa undergo very little HDX throughout the experiment, consistent with either hydrogen bonding of secondary and tertiary structure or a hydrophobic pocket. Of note, HDX decreases for the bound state in these peptides only after 1 h. The low initial HDX limits the dynamic range of binding-induced protection from HDX, but statistically significant protection is still observed.

**Figure 4.**
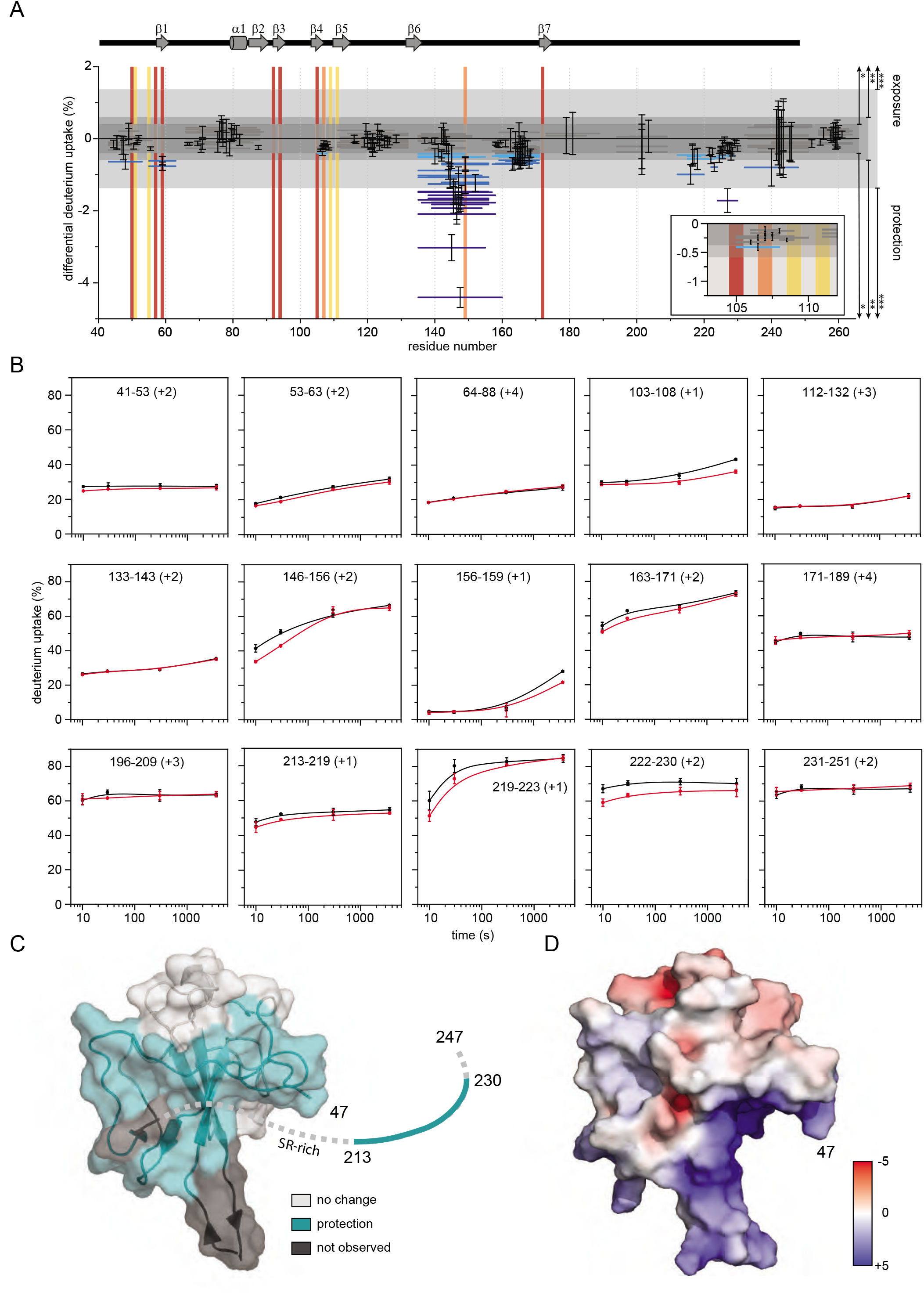
HDX-MS mapping of RNA binding to N_NTD-LKR S176D/S188D/S206D_. **A**. Woods’ plot showing cumulative differential HDX and validating differences using global significance limits. The horizontal bars depict the cumulative HDX differences between the RNA-bound and unbound N_NTD-LKR S176D/S188D/S206D_. Standard deviations are shown for each peptide. Peptides showing statistically significant differences are differentiated by global significance limit (*, p < 0.1; **, p < 0.05; ***, p < 0.01). The blue shade of the peptide bar indicates differing statistical significance (light blue, medium blue, and navy, respectively); gray peptide bars depict peptides where statistically significant differences in HDX were not observed. Vertical bars show previously reported binding sites (residues reported for RNA-binding CoV2 N-protein (Dinesh et al., 2020; Ye et al., 2020), AMP-binding HCoV-OC43 (Lin et al., 2014; Ye et al., 2020), and for both are shown in red, yellow, and orange, respectively). Secondary structure (PDB 6M3M) is shown above. **B**. Representative kinetic plots showing peptide level HDX as a function of exchange time (unbound, black; bound to RNA, red). **C**. Sites of protection measured by HDX mapped on the N_NTD_ structure (PDB 6M3M). Statistically significant HDX protection, regions of no difference in HDX, and regions where lacking proteolytic coverage results in no data are shown in teal, light gray, and dark gray, respectively. Those residues unresolved in the structure are shown as a dashed line, with the exception of those reporting a statistically significant difference in teal. **D**. Electrostatic potential calculated with APBS mapped on to the N_NTD_ structure (PDB 6M3M) shows a major positive charge groove. Red and blue represent negative and positive electrostatic potential. The color scale is in kT/e units.

Further, HDX analysis revealed that the protected regions (**Figure 4C**) overlap well with a basic patch groove in the N_NTD_ structure (**Figure 4D**). In addition, a region (213-230 aa) within the LKR domain, after the structured domain and SR-motif, shows statistically significant HDX protection. Interestingly, we did not detect HDX protection in the SR-motif, which was proposed to bind RNA. This may be due to the Ser-to-Asp mutations introduced into this region, changing the RNA binding patterns. Altogether, HDX results along with our biochemical data define an RNA binding interface within the NTD-LKR region.

### N_CTD_ is a sensitive serological marker

Current sero-diagnostic assays to identify COVID-19 positive individuals are based on the detection of antibodies against N due to its abundant expression and corresponding high immune response (Chew et al., 2020; Tang et al., 2020a, b). However, these N-directed serological assays are highly variable and their sensitivity depend on the sampling time-points, ranging from 0% to 93.75% (Liu et al., 2020; Tang et al., 2020a, b), suggesting that serological markers for SARS-CoV-2 infection can be further improved.

Given that the domains of N impact the various essential properties of N, we next assessed which domains contribute to the observed N immunodominance in RT-PCR confirmed COVID-19 patients. Plasma samples were collected from two cohorts, one in St. Louis, USA (n = 45) and one in Hong Kong (n = 23), at different time points of infection (Supplementary Table 1). Using these samples, we performed enzyme-linked immunosorbent assays (ELISAs) to detect IgG present in COVID-19 patient plasma to different N domains. First, we confirmed that purified N_WT_ is a sensitive serological marker to differentiate between COVID-19 positive and negative individuals (**Figure 5A**). As shown in **Figure 5B**, antibodies against all five N constructs were detected in the COVID-19 cohort (p < 0.0001 versus negative controls for all). A cut-off based on the mean of the negatives plus three standard deviations allowed us to assess the performance of each N construct at detecting IgG antibodies in COVID-19 positive individuals (**Figure 5C**). We find that N_NTD-LKR-CTD-Carm_ shows the lowest sensitivity (41.2%), whereas the truncated N_NTD-LKR-CTD_ can detect more COVID-19 positive individuals (70.6%). Furthermore, N_CTD_ shows the highest combination of sensitivity (75%) and specificity (96.4%) over the other N constructs tested. This is demonstrated by the lowest cut-off score for the N_CTD_ for negative control samples, despite a comparable level of amino acid sequence conservation of the N_CTD_ (29-41%) to the N_NTD_ (32-48%) and N_LKR_ (28-42%) domains with common cold corona viruses (Figure 1B).

**Figure 5.**
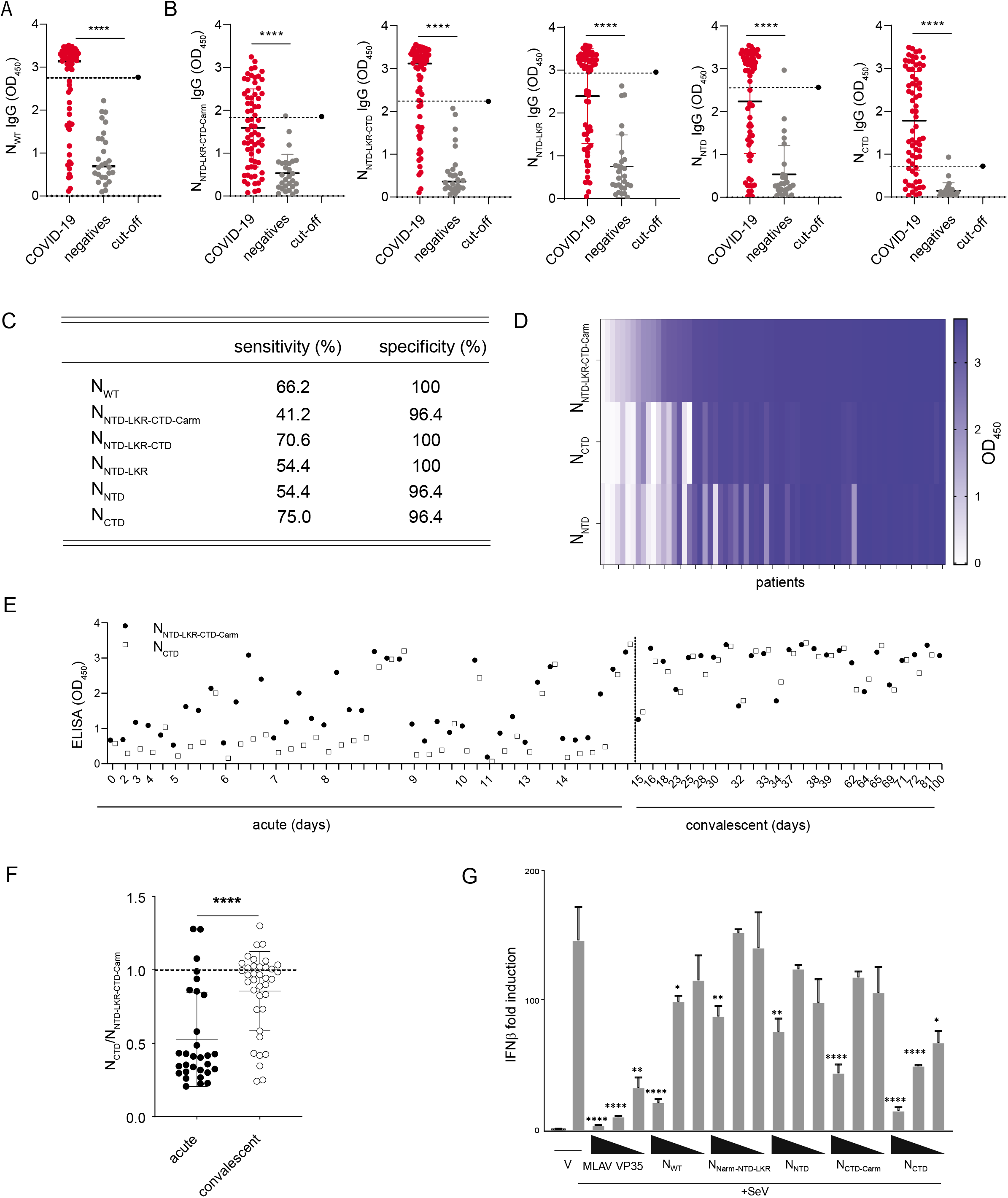
The CTD of N is a highly sensitive serological marker. **A.** ELISA data of N_WT_ screened against plasma of COVID-19 positive and negative individuals from a combined Hong Kong, PRC and St. Louis, MO, USA cohort. Black solid line indicates the mean OD_450_ value for each population. **** p < 0.0001. **B.** ELISAs with the various N constructs for patient IgG. ELISAs were performed on plasma samples from COVID-19 patients (n = 68) and negative controls (n = 28). The cut-off is represented by the dotted line and calculated as the mean + 3 standard deviations of the negative population. Mean values ± standard deviation of COVID-19 and negative groups are shown. **C.** Sensitivity and specificity for each of the N domains calculated from the ELISA results. **D.** Heat-map of ELISA results for N_NTD-LKR-CTD-Carm_, N_CTD_, and N_NTD_ constructs from COVID-19 samples (n = 67). Each column represents an individual sample. **E.** Maturation of the N_CTD_ and N_NTD-LKR-CTD-Carm_ IgG response over time (n = 67). **F.** Ratio of OD450 for N_CTD_ and N_NTD-LKR-CRD-Carm_ for acute and convalescent time-points. Mean values ± standard deviation of acute and convalescent COVID-19 samples are shown. Experiments were repeated twice. Statistical significance was calculated by unpaired Student’s t-test, ****p < 0.0001. **G**. Inhibition of SeV-induced IFNβ promoter activation by N constructs. Fold changes are relative to vector-only (V) transfections without SeV infection. MLAV VP35 served as a positive control for inhibition. Three transfection concentrations were used: 1.25, 12.5, and 125 ng/well. Statistical significance was determined by performing a one-way ANOVA followed with Tukey multiple comparison as compared to Sendai virus-infected control; **** p < 0.0001, *** p < 0.0002, ** p < 0.0021, * p < 0.0332.

We next compared the immunogenicity of N_NTD-LKR-CTD-Carm_ to N_CTD_ and N_NTD_ on an IgG heatmap during natural infection to an independent panel of 67 COVID-19 samples from Hong Kong. The magnitude of the IgG response to the N_NTD-LKR-CTD-Carm_ tends to follow the same trend as that of N_CTD_ (**Figure 5D**). When we assessed the kinetics of N_CTD_ and the N_NTD-LKR-CTD-Carm_ responses, we find that the magnitude of the N_CTD_ IgG detection tends to reach a similar level to that of N_NTD-LKR-CTD-Carm_ at convalescent time-points (after day 14) (**Figure 5E**). The ELISA ratio of N_CTD/NNTD-LKR-CTD-Carm_ demonstrates this finding, pointing to a maturation of the humoral immune response towards the N_CTD_ with time after infection (**Figure 5F**, p < 0.0001 for acute versus convalescent time-points). The early dominance of the N_NTD-LKR-CTD-Carm_ IgG response may reflect the recruitment of a crossreactive pre-existing N-specific response. This response becomes more specific with time for the N_CTD_ domain as a *de novo* antibody response is made.

Given that some RNA viral N proteins are known immune antagonists, including prior studies of SARS-CoV, we hypothesized that N from SARS-CoV-2 may also suppress the type-I interferon (IFN) signaling pathway (Messaoudi et al., 2015). Using an IFN-β promoter assay, we showed that N has a role in suppressing IFN signaling pathway when stimulated by Sendai virus (SeV) infection (**Figure 5G**). N_WT_ can inhibit IFN-β promoter activity, although not as well as Měnglà virus (MLAV) VP35, a potent inhibitor of IFN signaling (Williams et al., 2020). Both N_Narm-NTD-LKR_ and N_NTD_ show modest inhibition at the highest concentration tested. However, N_CTD-Carm_ shows similar levels of inhibition as N_WT_, and N_CTD_ displays the highest inhibition even at lower protein concentrations. In summary, N is a potent inhibitor of IFN signaling and the CTD domain appears to be the region critical for mediating this function.

## Discussion

SARS-CoV-2 N protein is a core viral protein produced by the subgenomic RNA, positioned proximal to the 3’ end of genome, display high transcription levels, and is in high abundance in virions. N is prone to forming higher-order oligomers that have a role in binding to the RNA genome of SARS-CoV-2 and nucleocapsid formation. N is also regulated by post-translational modifications, including phosphorylation, which changes the physiochemical properties of N and likely directs its multiple roles at different stages of the viral replication cycle.

Here we used a series of N constructs to dissect how each domain contributes to oligomerization and RNA binding, and how phosphorylation can modulate these properties. N oligomerization and RNA binding are likely linked; N oligomerization provides a platform for high affinity RNA interactions whereas genomic RNA serves as a string connecting N proteins. The modular domains of N provide multiple regulatory layers for genomic access. For example, phosphorylation of N in the LKR region reduces RNA binding to N and modify distribution of N-RNA species.

We also gained insight into the antigenicity of individual domains of N and its potential utility in serological studies. We data revealed that N_CTD_ acts more specifically in detecting infection of SARS-CoV-2, the causative agent of COVID-19, from plasma in comparison to N_WT_. Consistent with our observation, the N_CTD_ region is also predicted to encompass major antigenic sites of N (Bussmann et al., 2006; Liang et al., 2005). Owing to the relative conservation of N within coronaviruses, it is crucial to understand how N-directed antibodies generated by different coronaviruses are cross-reactive with those that are derived when exposed to SARS-CoV-2. The common cold coronavirus protective immunity is short-lasting (Edridge et al., 2020); reinfection with the same seasonal coronavirus occurred frequently at 12 months after the initial infection using an ELISA approach against the N C-terminal region (coronavirus NL63, 215-377 aa; 229E, 213-389 aa; OC43, 328-428 aa; HKU1, 326-441 aa), which is similar to the N_CTD_ region we used (SARS-CoV-2, 248-369 aa). Interestingly, 2 out of 10 individuals assessed in this study of longitudinal donors unexposed to SARS-CoV-2 by Edridge *et al.,* produced broadly reactive antibodies towards SARS-CoV-2 N_WT_. The possibility of broadly reactive antibodies in unexposed individuals highlights the need for domain specific serology, such as our use of the N_CTD_ for increased sensitivity to discriminate COVID-19 cases, while reducing the false-positive rate from cross reactive antibodies generated by infections of the common cold coronaviruses.

In conclusion, we describe our efforts to characterize domain specific insights into essential biochemical and serological properties of SARS-CoV-2 N. Our results advance the understanding of viral replication processes and highlight the diagnostic value of using N domains as a highly specific and sensitive markers of COVID-19. While our study highlights the important functions associated with N_CTD_, much remains to be characterized, including the mechanistic link between the high immunogenicity of N_CTD_ and the physical properties such as RNA binding and oligomerization.

## Methods

### Patients and sample collection

Our study enrolled a total of 67 patients with RT-PCR confirmed COVID-19 infection: with 45 patients from St. Louis, MO, USA, and 23 patients from Hong Kong, PRC. The negative samples (n = 28) used in this study were from St. Louis, USA, and were obtained from patients following the start of the pandemic. Plasma samples were obtained from patients at Barnes-Jewish Hospital (St. Louis, MO, USA) and the Hong Kong Island West Cluster of Hospitals (Hong Kong, PRC). Both hospital systems are urban, tertiary-care, academic medical centers. Positive and negative patients from all cohorts were confirmed using standard of care, RT-PCR based methods. The collection of patient plasma was approved by the Human Research Protection Office at Washington University in St. Louis (IRB reference number 202007097) and the Institutional Review Board of The Hong Kong University and the Hong Kong Island West Cluster of Hospitals (IRB reference number UW20-169). Plasma samples were collected from heparinized blood. Sample day was defined as days post-symptom onset.

### Enzyme-linked immunosorbent assay (ELISA)

ELISA assays were performed with Nucleoprotein (N) proteins made in house, as described below. Briefly, recombinant N proteins were coated on 96 well flatbottom immunosorbent plates (Nunc Immuno MaxiSorp) at a concentration of 500 ng/mL, in 100 μL coating buffer (PBS with 53% Na_2_CO_3_ and 42% NaHCO_3_, pH 9.6) at 4°C overnight. An additional plate coated with a non – specific protein (blocking buffer, PBS with 5% fetal bovine serum (FBS)) was used to measure the background binding of each plasma sample. Following FBS blocking and thorough washing, diluted plasma samples (1:100) were bound for 2 hours, further washed and then detected by an anti – human IgG secondary antibody labelled with HRP (Invitrogen), and absorbance detected at 450nm on a spectrophotometer (Wallac).

### Protein Expression and Purification

SARS-CoV-2 N constructs were expressed as His-tag fusion proteins in BL21 (DE3) *E. coli* cells (Novagen). At OD_600_ of 0.6-0.7, recombinant protein expression was induced with 0.5 mM isopropyl β-d-1-thiogalactopyranoside (IPTG) for 12-14 h at 18°C. Cells were harvested and resuspended in lysis buffer containing 20 mM Tris (pH 7.5), 1 M NaCl, 20 mM imidazole, 5 mM 2-mecaptoethanol (BME). Cells were lysed using an EmulsiFlex-C5 homogenizer (Avestin) and lysates were clarified by centrifugation at 30,000 × *g* at 4 °C for 40 min. N proteins were purified using affinity tag and gel filtration columns. Purity of N proteins were determined by Coomassie staining of SDS-PAGE.

### Negative Staining EM

2 μL of N sample at a concentration of 1 mg/mL was applied to a glow-discharged copper grid (Ted Pella), washed twice with water before staining with 2% uranyl acetate for 30 s and air dried. Grids were imaged using a JEOL JEM-1400plus Transmission Electron Microscope operating at 120 kV and recorded with an AMT XR111 high-speed 4k × 2k pixel phosphor-scintillated 12-bit CCD camera.

### Dynamic Light Scattering (DLS)

DLS experiments were performed on a DynaPro-PlateReader II (Wyatt Technologies Corporation). Measurements of N samples in triplicates (1 mg/mL) were obtained at 25 °C and analyzed using Dynamics software (Wyatt).

### Fluorescence Polarization Assay (FPA)

FPA experiments were performed on a Cytation5 plate reader (BioTek) operating on Gen5 software. Excitation and emission wavelengths were set to 485 and 528 nm, respectively, with a bandpass of 20 nm. Read height and G factor were set to 8.5 mm and 1.26, respectively using the autogain function. For RNA binding experiments, fluorescein isothiocyanate (FITC) labelled 20 nt-ssRNA or 19 nt slRNA at a final concentration of 1 nM was loaded on N samples (in 20 mM HEPES (pH 7.5), 150 mM NaCl, 2 mM TCEP, 5% glycerol) at concentrations ranging from 0.4 nM to 10 μM in a 96-well plate. After 10 min of incubation, fluorescence polarization signals were read. The fluorescence polarization values were then plotted against N concentrations to fit the dissociation constant, *K_D_*, using ORIGIN software. For anisotropy plots, anisotropy values were converted from polarization according to previous research (Kozlov et al., 2012).

### LC-MS Analysis

Unless otherwise indicated, all chemical reagents were sourced from Millipore Sigma and used without further purification. For LC-MS analyses, 30 pmol of protein in 50 μL of 1:1 solvent mixture of acetonitrile:water with 0.1% formic acid (CovaChem) was loaded onto a C8 trap (ZORBAX Eclipse XDB C8 column, 2.1 × 15 mm, Agilent), desalted for 3 min by using water/0.1% formic acid at a flow rate of 100 μL/min, and eluted using an 14 minute gradient from 0 to 80% acetonitrile/0.1% formic acid at a flow rate of 100 μL/min. Samples were analyzed using a MaXis 4G Q-TOF (Bruker Daltonics). The mass spectrum was extracted guided by the elution peak and submitted to PMI Intact Mass and searched for M values ranging from 5-50 kDa.

### HDX-MS

N_NTD-LKR S176D/S188D/S206D_ was incubated with a 20-nt ssRNA at a 1:1 ratio. After incubation, 2 μL of 50 μM protein/protein-RNA in PBS (pH 7.4) was diluted 10-fold (v/v) with labeling buffer (PBS in D_2_O, pD 7.0) (D_2_O from Cambridge Isotope Laboratories), incubated for 10, 30, 300, and 3600 s on ice, quenched by using a 60% dilution with 3 M urea, PBS (pH 2.5), and flash frozen for later LC-MS analysis. A 0 s control was prepared with PBS in H_2_O. Prior to incubation, each 50 μL of 2 μM sample was thawed for 1 min at 37 °C before injection into a custom-built liquid chromatography (LC) apparatus for LC-MS analysis. The labelled protein passed through two in-house packed protease columns (2 mm × 20 mm), coupled so that the first using protease from *Aspergillus saitoi* type XIII (FXIII) and the second porcine pepsin (0.1% formic acid, flow rate 200 μL/min); the resulting peptides were trapped on a ZORBAX Eclipse XDB C8 column (2.1 mm × 15 mm, Agilent), desalted for 3 min, and then separated on a Hypersil Gold C18 column (2.1 × 50 mm, Thermo Fisher) with a 10.5 min linear gradient from 4 – 40% acetonitrile/0.1% formic acid (flow rate 100 μL/min). All valves, tubes, and columns (except for the protease columns, which lose activity at low temperature) were submerged in ice during the experiment to minimize back exchange. Peptides were eluted into a Bruker Maxis HM Q-TOF MS for mass analysis. Experiments were in duplicate unless otherwise indicated. The HDX data processing was performed by using HDExaminer (version 2.5.1, Sierra Analytics, Inc.).

### IFN-β promoter reporter gene assay

HEK-293T cells (5 × 10^4^) were co-transfected using Lipofectamine 2000 with 25 ng of an IFN-β promoter-firefly luciferase reporter plasmid, 25 ng of pRL-TK *Renilla* luciferase reporter plasmid, and 125, 12.5, and 1.25 ng of the indicated viral protein expression plasmid. Twenty-four hours post-transfection, cells were mock-treated or SeV (15 hemagglutination units / ml) infected. Eighteen hours post-treatment or post-infection, cells were lysed and analyzed for luciferase activity using a Dual-Luciferase reporter assay system (Promega). Firefly luciferase activity was normalized to *Renilla* luciferase activity. Assays were performed in triplicate; error bars indicate the standard error of the mean (SEM) for the triplicate. Viral protein expression was confirmed by Western blot analysis.

## Acknowledgements

We thank Dr. N. Krogan (UCSF) for sharing SARS-CoV-2 plasmids, Drs. A. Holehouse and A. Soranno (Washington University School of Medicine) for providing critical feedback, and R. Ridings for coordinating studies between WUSM and external groups. Research was supported by Fast Grant ##2161 (Emergent Ventures) to G.K.A. and NIH grants (P01AI120943, R01AI123926 to C.F.B, G.K.A. and C.F.B; R01AI107056 to D.W.L.; P41GM103422 and R24GM136766 to M.L.G.; P01AI120943, R01AI143292 and R01AI148663 C.F.B). S.A.V was supported by COVID190115 and COVID190126 Health and Medical Research Fund, Food and Health Bureau, Hong Kong and NIH/NIAID CEIRS contract HHSN272201400006C. A.J.Q. is supported by a NIH T32 training grant (T32CA009547).

## Author contributions

CW, GKA, and DWL conceived the overall project. All authors were integral to the design and execution of the study. CW, AQ, GKA, and DWL wrote the initial draft with significant input from all authors.

## Supplementary Figure Legends

**Supplementary Figure 1, related to Figure 1.**
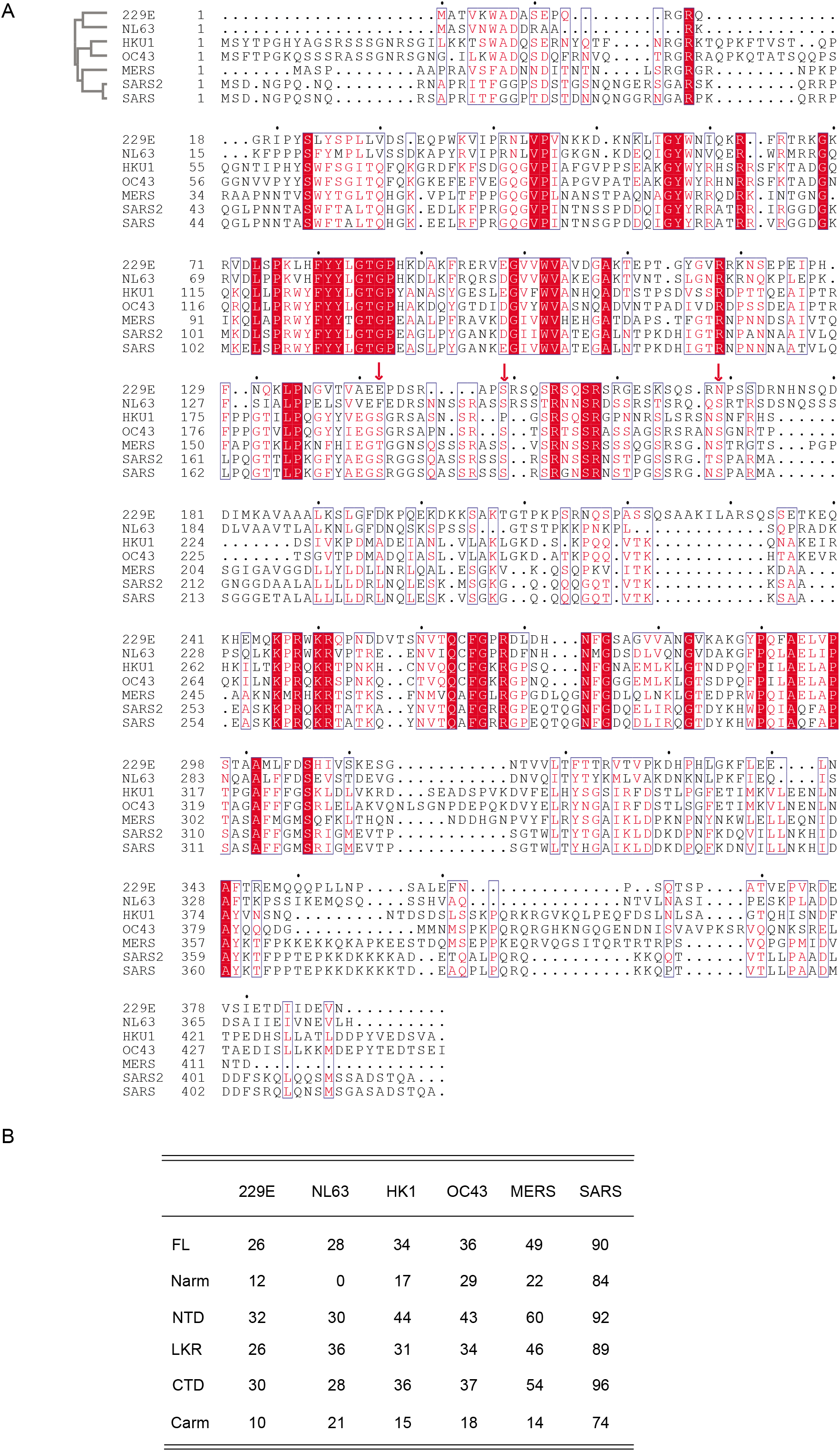
Multiple sequence alignment of coronavirus nucleocapsids. **A.** Multiple sequence alignment of coronavirus nucleocapsids. Sequences were aligned using Clustal Omega. Accession numbers used are 229E (APT69891.1), NL63 (YP_003771.1), HK1 (AAT98585.1), OC43 (AAR01019.1), MERS (AKL80590.1), SARS (AAP30037.1), SARS2 (YP_009724397.2). Alignments were analyzed using ESPript3. The three serines (176, 188, and 206) are labeled with red arrows. **B**. Sequence identity between SARS-CoV-2 N and that of common cold coronaviruses and MERS and SARS. FL, full length. All units are in %. Percent identity matrixes for corresponding domains of N are generated using Clustal2.1.

**Supplementary Figure 2, related to Figure 1.**
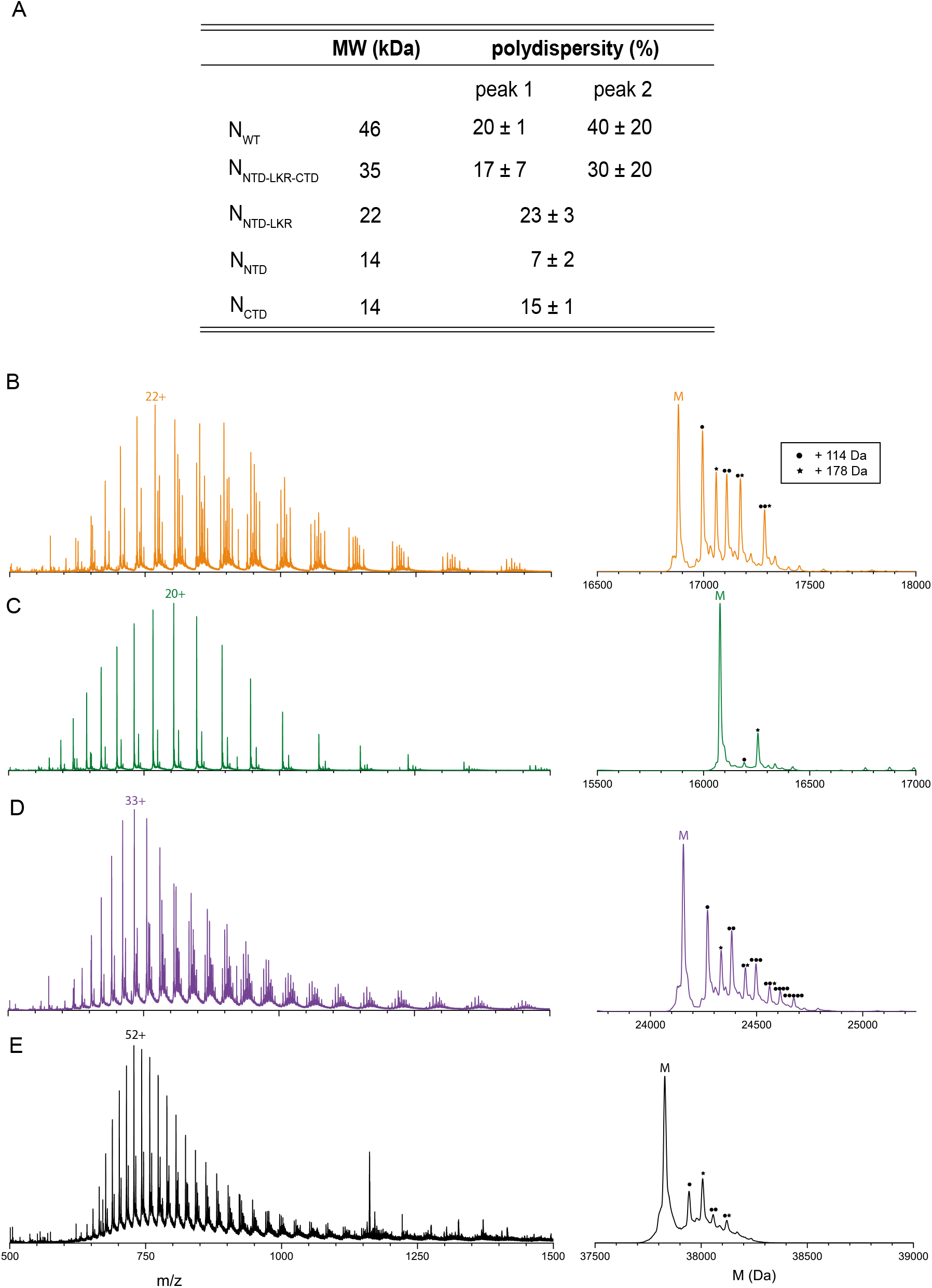
Denaturing mass spectra of N protein truncations N_NTD, NCTD, NNTD-LKR_, and N_NTD-LKR-CTD_. **A.** DLS polydispersity table for N constructs. Higher values indicate broader size distributions. Numbers are reported as average and standard deviation of three experiments. Deconvolution yields experimental M values of 16,881 Da, 16,078 Da, 24,155 Da, and 37,829 Da for (**B**) N_NTD_, (**C**) N_CTD_, (**D**) N_NTD-LKR_, and (**E**) N_NTD-LKR-CTD_ respectively, matching theoretical values within 1 Da, based on protein sequence. Deconvoluted mass spectra (right) and adduct series corresponding to pervasive trifluoroacetic adducts (delta mass 114 Da, circle) and a-N-gluconoylation (delta mass 178 Da, star). TFA adducts are introduced by the ion pairing reagent in solvent, while α-N-gluconoylation is a common modification occurring on His-tagged proteins. Native spray of NTD (not pictured) yielded no peaks with delta mass 114 Da, but retained a single delta mass 178 Da, confirming transient TFA adducts are an artifact of the denaturing experiment, but the α-N-gluconoylation of the His-tag is covalent.

**Supplementary Figure 3, related to Figure 2.**
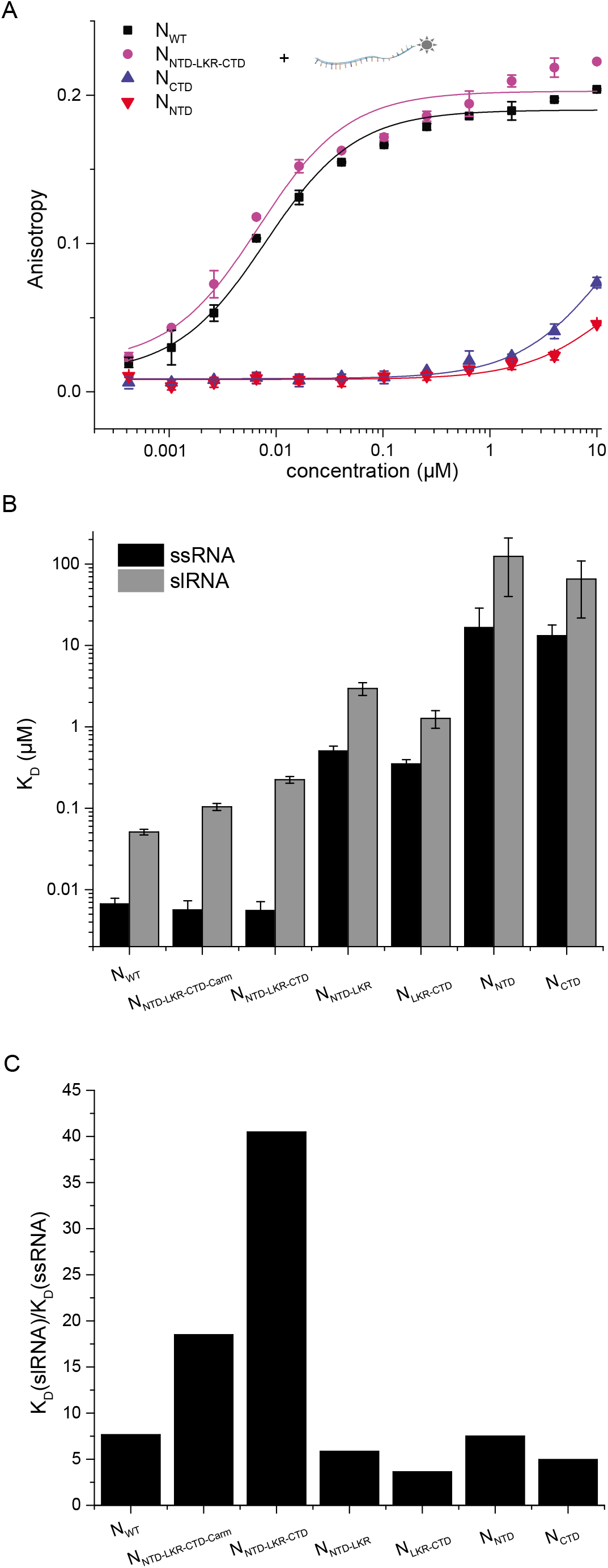
Nucleocapsid binds stem-loop RNA with reduced affinity. **A.** Fluorescence anisotropy binding curves of N constructs to a 20-nt ssRNA. Anisotropy values were converted from polarization according to previous research (Kozlov et al., 2012). The fitted K_D_ values are 0.007 ± 0.001 μM (N_WT_, black square), 0.006 ± 0.002 μM (N_NTD-LKR-CTD_, magenta circle), 14 ± 5 μM (N_CTD_, blue up triangle) and 18 ± 14 μM (N_NTD_, red down triangle). These values are very close to those of polarization. In this system, binding monitored by anisotropy is similar to that of polarization. **B.** Fitted KD values for N constructs binding to ssRNA (black) and slRNA (grey). **C.** Ratio of K_D_ of slRNA over that of ssRNA for N constructs. The reduced binding to slRNA is around 5-fold for most N constructs. The reduction is higher for those of N_NTD-LKR-CTD-Carm_ and N_NTD-LKR-CTD_, suggesting Narm and Carm are more involved in slRNA binding.

**Supplementary Figure 4, related to Figure 3.**
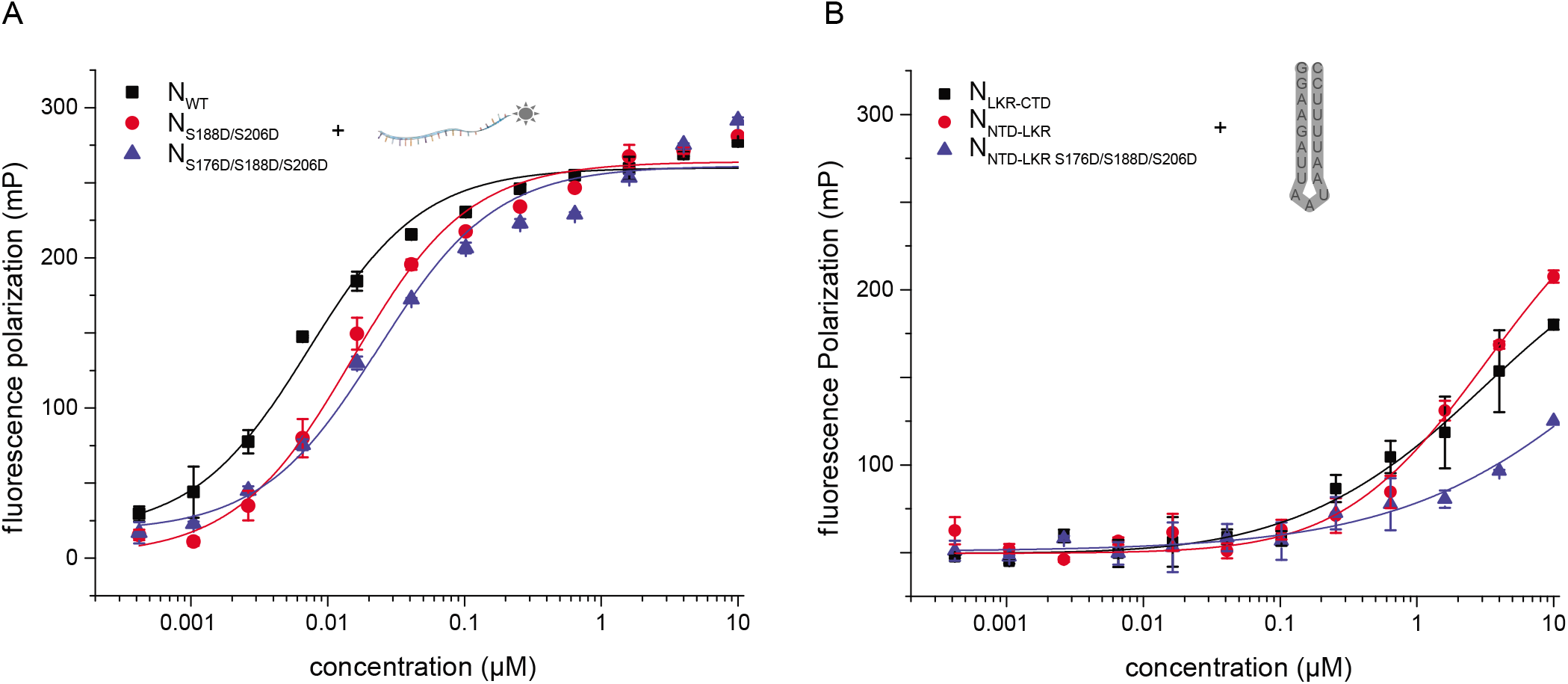
Phosphorylation mimetics of N reduce RNA binding. **A.** Fluorescence polarization binding curves of N constructs to a 20-nt ssRNA. The fitted K_D_ values are 0.007 ± 0.001 μM for N_WT_, 0.015 ± 0.002 μM for N_S188D/S206D_, and 0.023 ± 0.006 μM for N_S176D/S188D/S188D_. **B.** Fluorescence polarization binding curves of N constructs to a 19-nt slRNA. The fitted K_D_ values are 1.3 ± 0.3 μM (N_LKR-CTD_, black square), 3.0 ± 0.5 μM (N_NTD-LKR_, red circle), and 2.9 ± 1.4 μM (N_NTD-LKR_ S176D/S188D/S206D, blue up triangle).

**Supplementary Figure 5, related to Figure 4.**
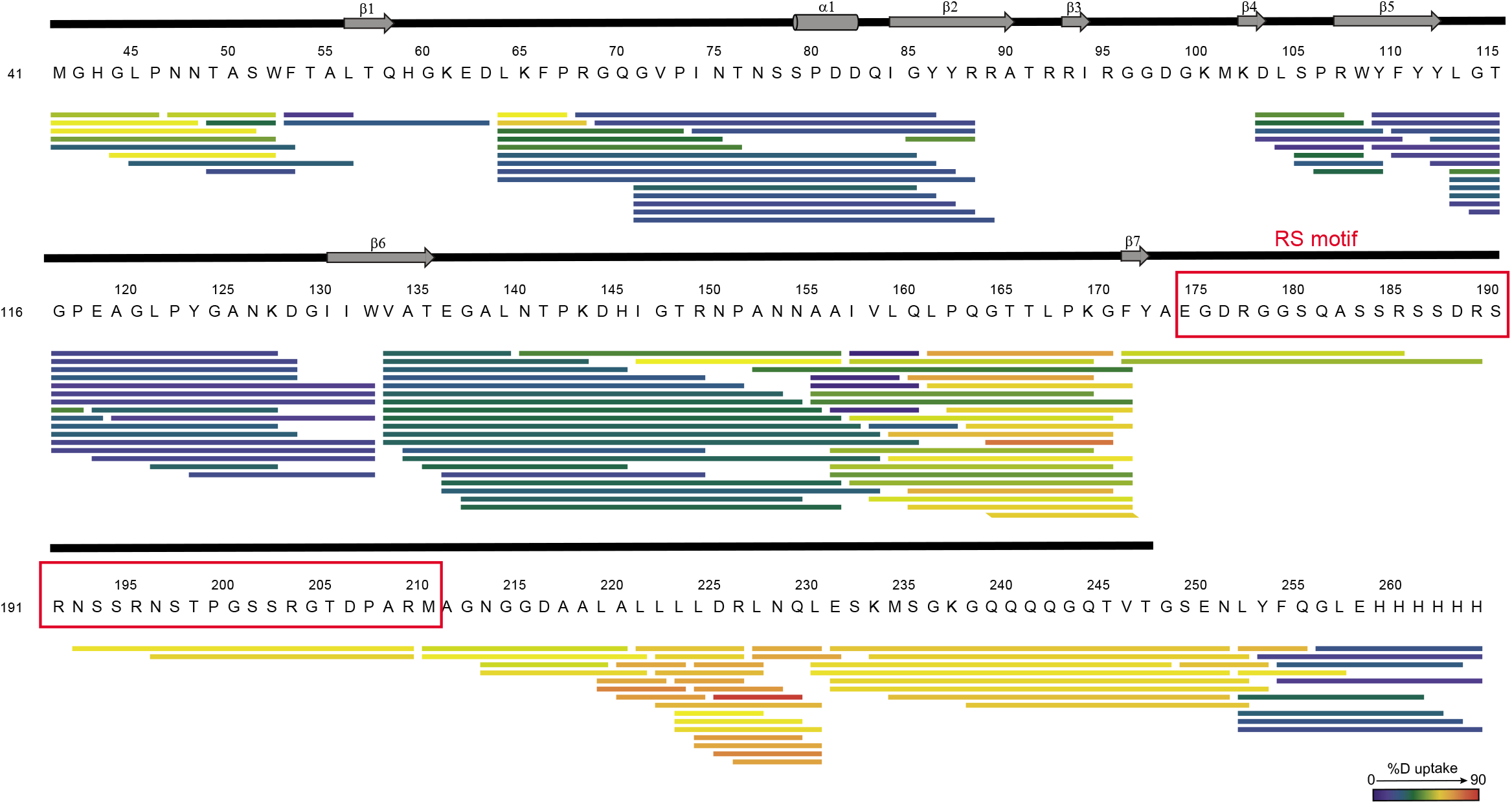
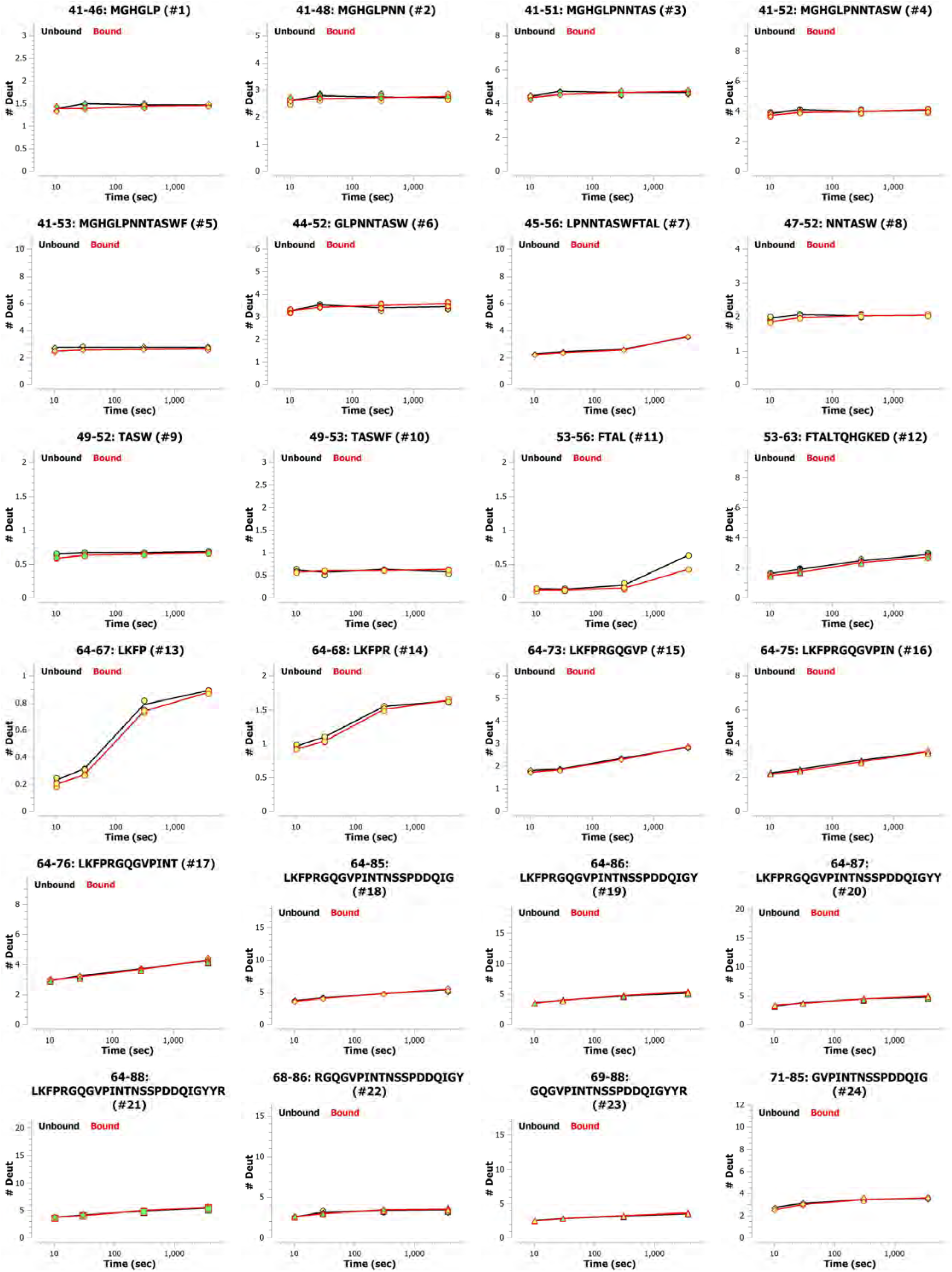
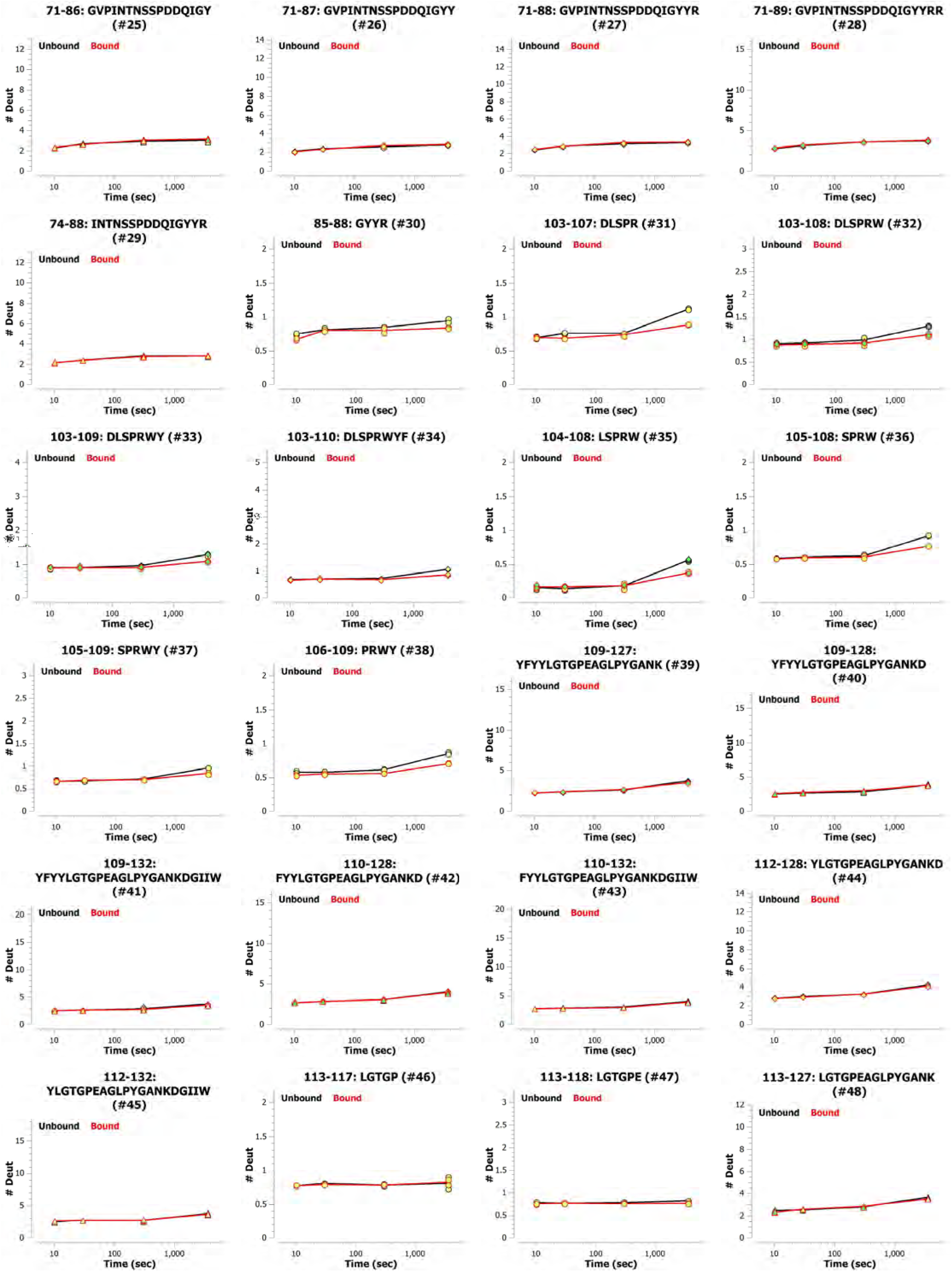
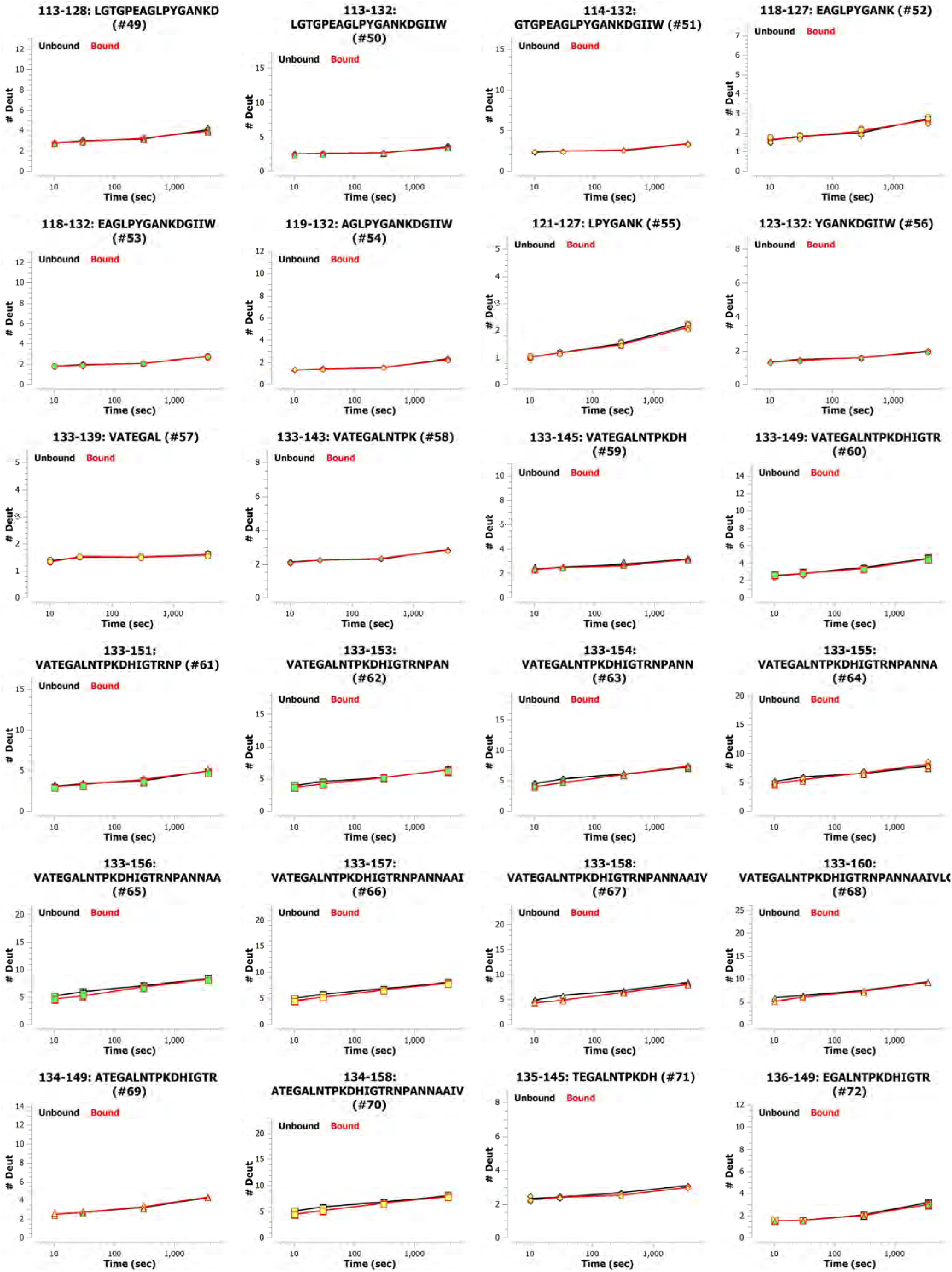
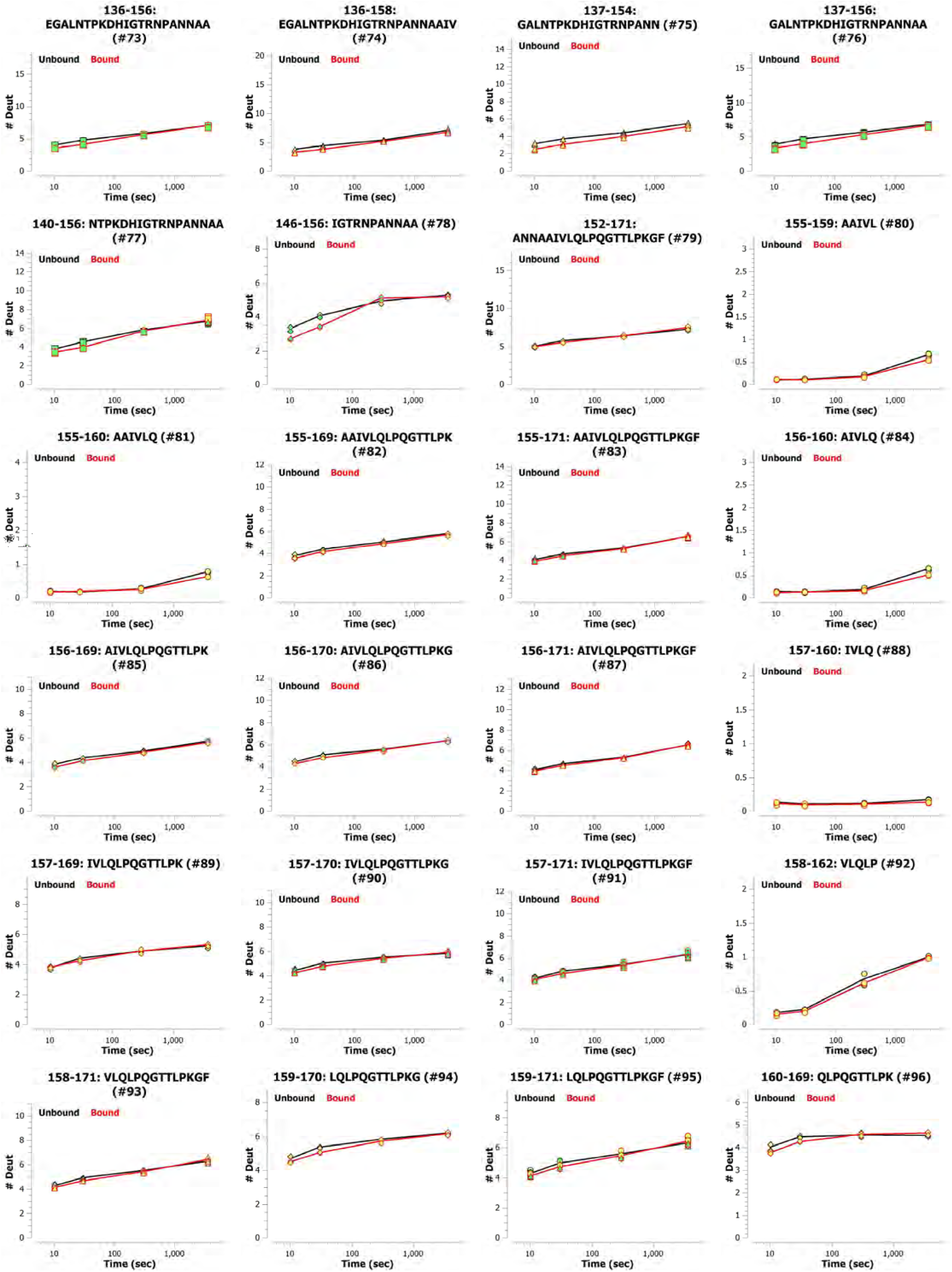
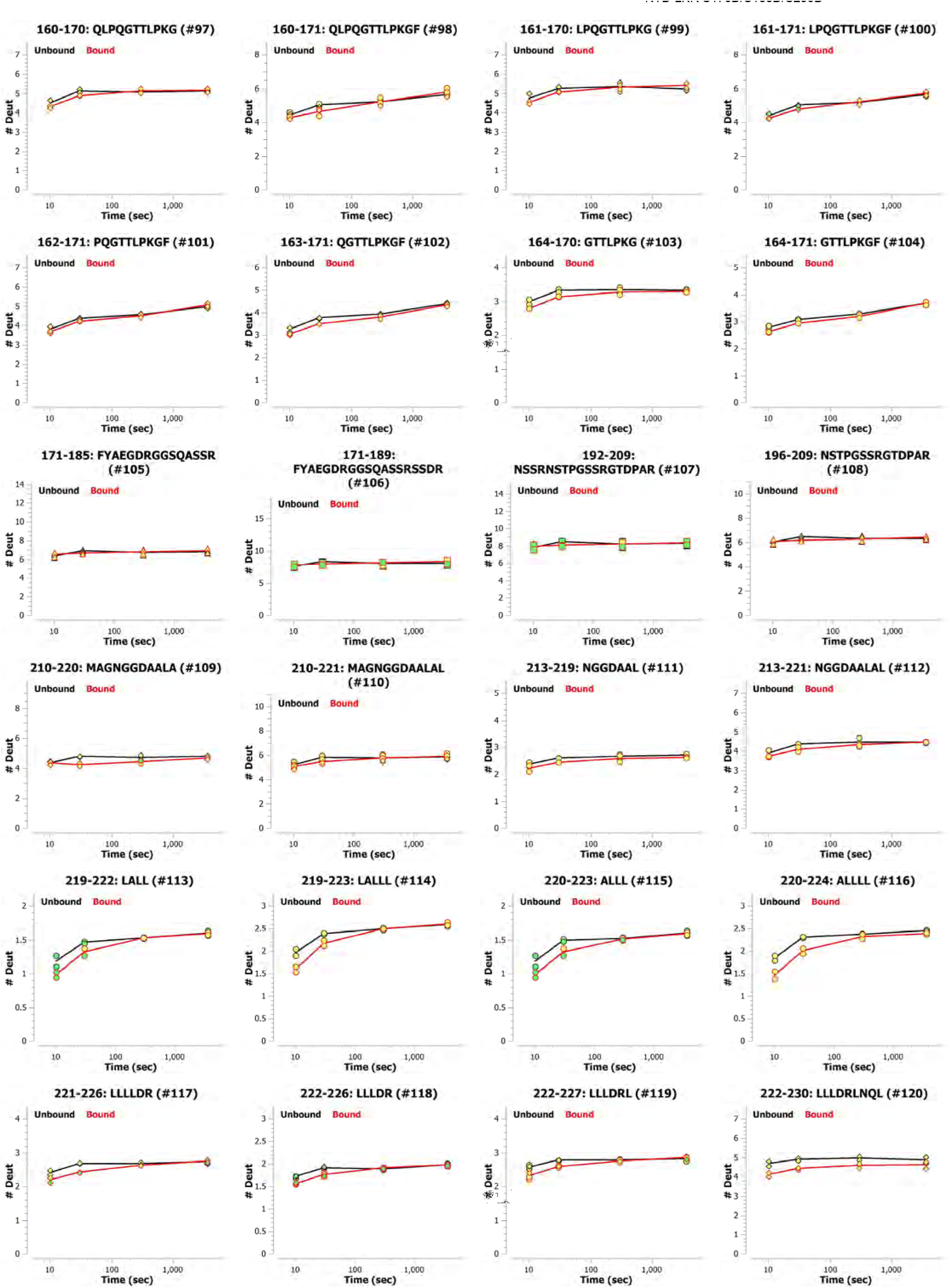
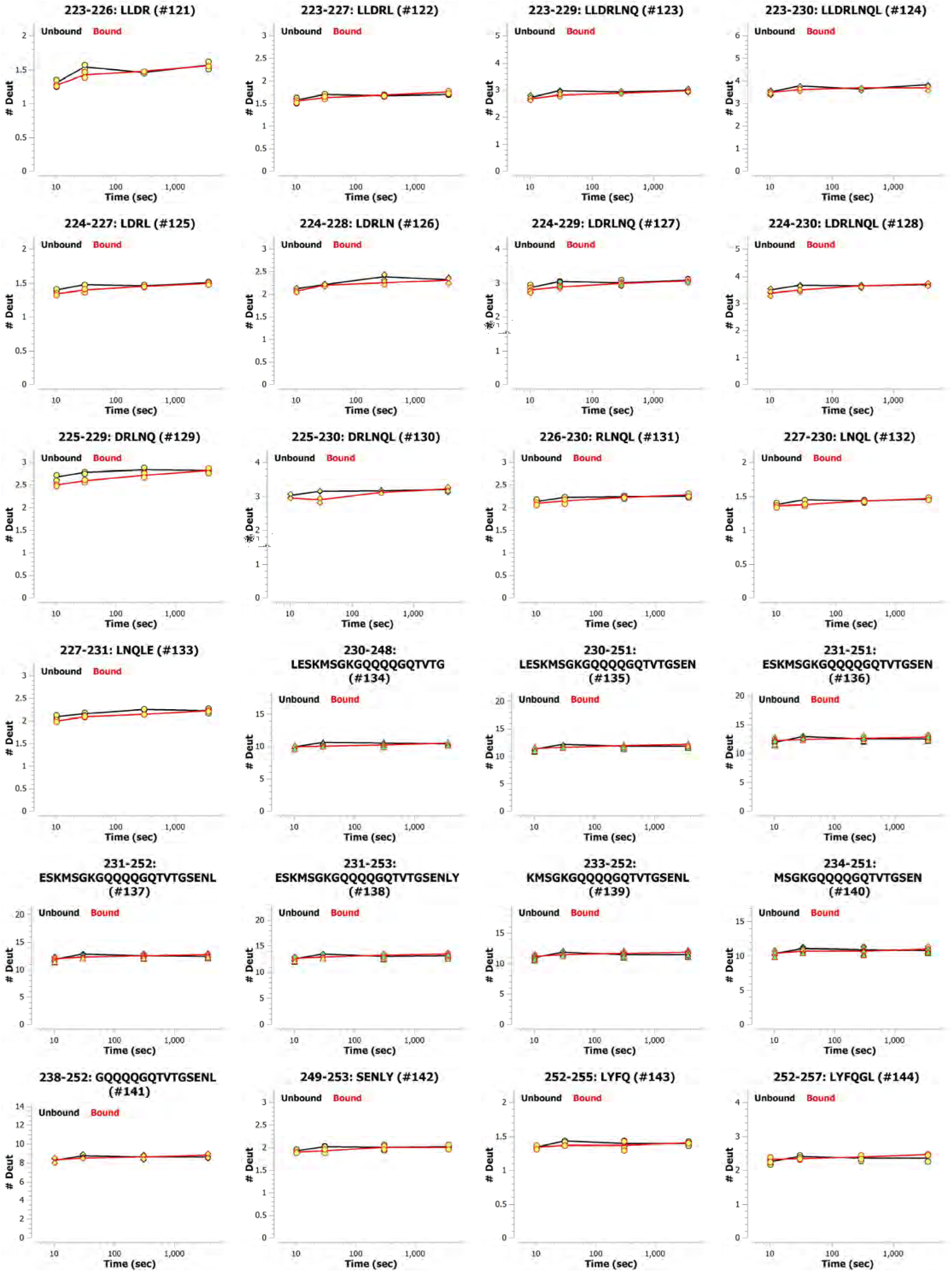
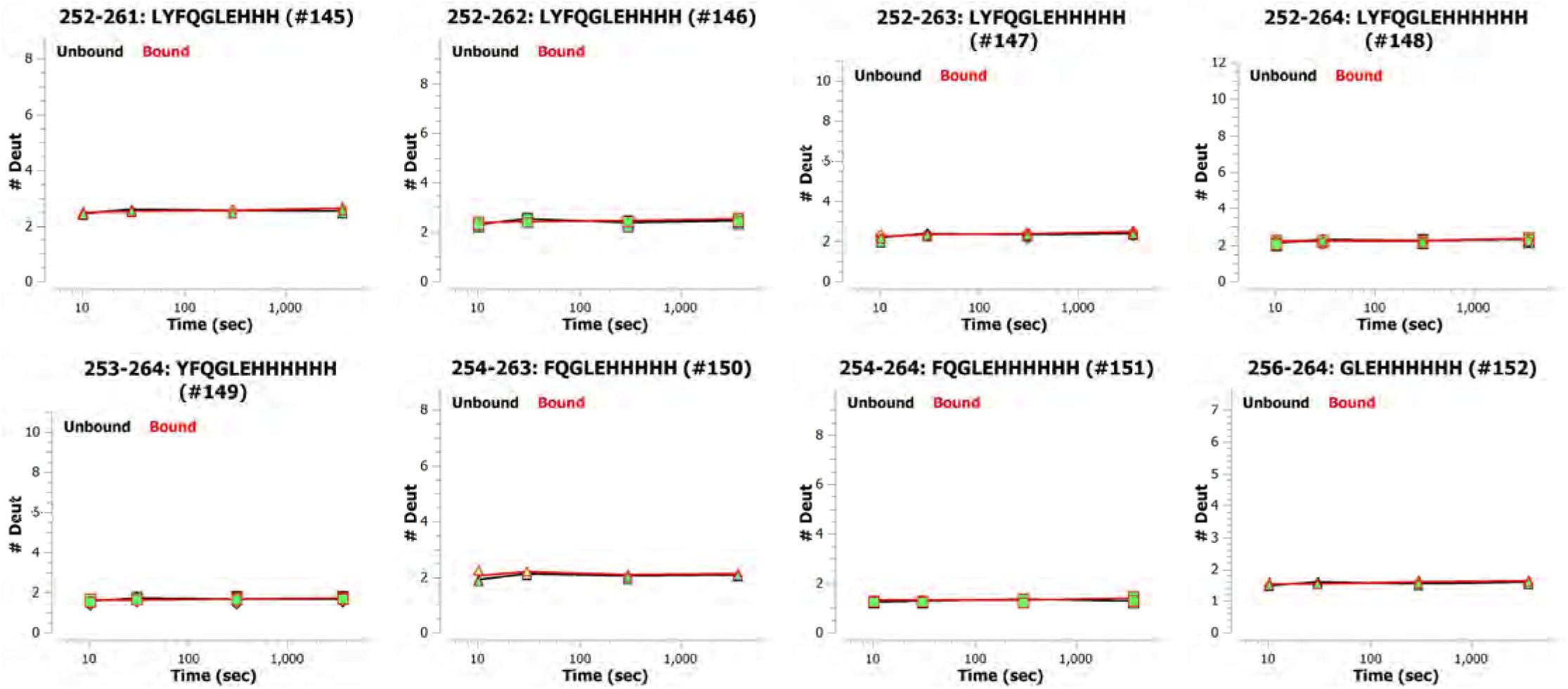
Sequence coverage of N_NTD-LKR S176D/S188D/S206D_ in HDX-MS and HDX of the unbound state. **A**. Protein coverage map of unbound state N_NTD-LKR S176D/S188D/S206D_ HDX yielding 152 peptides with 93.3% sequence coverage. Peptide bars are colored according to their average %HDX relative to the color bar, where cooler colors depict low average %HDX and warmer colors depict high average %HDX. The secondary structure reported by PDB 6M3M is shown above the sequence. Overall, the HDX of the unbound state is largely consistent with the reported secondary structure and a well-ordered tertiary structure; regions outside of the reported structure undergo relatively rapid HDX, consistent with a lack of backbone hydrogen bonding. Interestingly, despite a lack of reported secondary structure in the region of 155-160, relatively low HDX was observed, consistent with either hydrogen bonding of secondary/tertiary structure or a hydrophobic pocket. SR-motif in LKR are boxed in red. **B**. All kinetic plots used in the peptide-level difference plot in Figure 5A show peptide level HDX as a function of exchange time (unbound, black; bound to RNA, red).

